# Large-scale AI-guided discovery of antifreeze peptides reveals functional tandem-repeat architectures

**DOI:** 10.64898/2026.11.03.763275

**Authors:** Yijun Li, Chi Wang, Jialin Liu, Mingye Song, Letian Zhang, Shuzhe Sun, Wentao Li, Jiangjie Qiu, Yihua Qin, Jinhao Yao, Leyi Zhao, Huimin Yu, Xiaonan Wang

## Abstract

Despite growing demand for biocompatible materials that regulate ice crystallization in cryopreservation, antifreeze peptide discovery remains constrained by limited candidate sequence diversity and reliance on known protein scaffolds. Here, we establish an AI-guided discovery framework integrating multimodal antifreeze protein prediction, generative sequence expansion and local sequence-pattern analysis. AFPFusionLM combines protein sequence and structure-aware representations, achieving an AUROC of 0.937 and an AUPRC of 0.957 on an independent reference set. Antifreeze-domain-adapted protein sequence generation and multi-model consensus screening of UniProt, glacier metagenomes and generated sequences yielded 87,305 candidate protein records. Large-scale sequence analysis revealed recurring periodic sequence architectures across multiple candidate protein clusters, enabling the identification of short-peptide families with shared repeat features and their natural variants. Experimental evaluation revealed ice recrystallization inhibition activity in multiple peptides from repeat-associated families, with inhibitory effects comparable to or greater than those of reference peptides reported in the literature. Cell-based experiments using a representative peptide-containing formulation also supported its cryoprotective potential under reduced-DMSO conditions. These findings show that large-scale sequence exploration can reveal function-associated peptide sequence architectures, providing a research framework for the systematic discovery and subsequent optimization of antifreeze peptides.

## Introduction

Cryopreservation is essential to cell therapy, regenerative medicine, biobanking and the long-term storage of biological products. However, ice formation, growth and recrystallization during freezing and warming can damage cell membranes, disrupt osmotic balance and impair cellular function, limiting preservation efficiency [1, 2]. Conventional permeating cryoprotectants, including dimethyl sulfoxide (DMSO), glycerol and ethylene glycol, are widely used, but their application remains constrained by concentration- and cell-type-dependent toxicity, osmotic stress and the need for subsequent removal [2–4]. Developing cryoprotective materials that combine effective control of ice with favourable biocompatibility is therefore an important objective. In nature, antifreeze proteins that have evolved in cold-adapted organisms inhibit ice growth and recrystallization through interactions with ice surfaces, providing a basis for bioinspired cryoprotective materials [5]. Nevertheless, the relatively high molecular weights and complex higher-order structures of some natural antifreeze proteins, together with challenges in isolation, purification and large-scale production and concerns about potential immunogenicity, limit their direct application [6]. Antifreeze peptides offer lower molecular weights, defined compositions and straightforward chemical synthesis and sequence optimization, making them attractive for developing structurally controllable ice-regulating materials with potential for favourable biocompatibility [6–10]. Efficient discovery of functionally active antifreeze peptides nevertheless remains a central challenge.

Antifreeze peptide research faces two interrelated bottlenecks: limited exploration of candidate sequence space and a dependence on existing sequence and structural priors. Conventional studies typically start from known antifreeze proteins, conserved ice-binding motifs or specific structural templates, construct candidate libraries through sequence truncation, amino acid substitution and modular assembly, and then use experiments and molecular simulations for screening and optimization [6–10]. Although this approach enables precise design within local sequence neighbourhoods, it cannot readily encompass the extensive diversity of natural proteins. Machine learning has recently been applied to antifreeze peptide identification and design, providing computational tools for candidate identification [11, 12]. However, broader diversification of candidate sources and coordinated exploration across multiple sequence resources remain limited, restricting systematic discovery beyond conventional annotations and established design templates. Antifreeze peptide discovery therefore needs to move beyond local optimization around a small number of sequences towards systematic exploration of broad protein sequence space.

Advances in protein language models and generative artificial intelligence provide a technological basis for this transition [13, 14]. The scientific value of large-scale sequence exploration lies not only in expanding candidate numbers but also in revealing functional sequence patterns that are difficult to detect in small-scale studies. By resolving the chemistry and spacing of ice-binding sites and their interactions with interfacial water, previous studies have connected specific molecular structures to antifreeze activity and established principles for rational peptide design [8, 9, 15]. These studies, however, primarily describe structure–function relationships within particular molecular scaffolds and their neighbouring sequence space, rather than the functional diversity of broader protein sequence space. Natural and generated proteins may contain as-yet-unrecognized local sequence organizations that confer antifreeze activity on short peptides. Systematic identification and functional evaluation of these patterns could advance the field from optimizing established structures towards discovering new functional architectures.

Here, we move beyond template-dependent design to establish an antifreeze peptide discovery framework in which large-scale sequence exploration drives the identification of functional patterns. The framework integrates AI-based prediction, generative expansion, sequence-pattern mining, molecular simulation and experimental validation. Systematic exploration of public protein databases, polar metagenomes and generated sequences yielded 87,305 potential antifreeze protein candidates retained by multimodel consensus screening. More importantly, expanding the search revealed biological features beyond an increase in candidate numbers. We identified periodic sequence architectures in multiple candidate protein clusters, allowing previously dispersed local features to emerge as recurring patterns with potential functional relevance. Experimental validation confirmed ice recrystallization inhibition by related repeat-containing peptides, while molecular simulations suggested that periodic spatial arrangements of hydrophilic and hydrophobic residues may promote antifreeze function by jointly modulating ice-interface interactions and local structural stability. This study expands the searchable space for antifreeze peptides and translates that expansion into functional sequence architectures and mechanistic leads, establishing a route towards systematic discovery and interpretable design.

## Results

### An AI-guided framework for antifreeze peptide discovery

To systematically explore potential antifreeze-active fragments across broad protein sequence space, we established an AI-guided framework that integrates protein function prediction, generative sequence expansion and local sequence-pattern discovery (Fig. 1a). The framework draws on three complementary sources: public protein databases, glacier metagenomes and artificially generated proteins. These sources provide natural protein diversity, incompletely annotated environmental sequences and additional candidate space created by generative modelling, respectively. A common prediction and screening workflow organizes proteins from these sources into a candidate collection for comparative analysis, providing a foundation for discovering functional short peptides within large protein sequence libraries.

**Figure 1.**
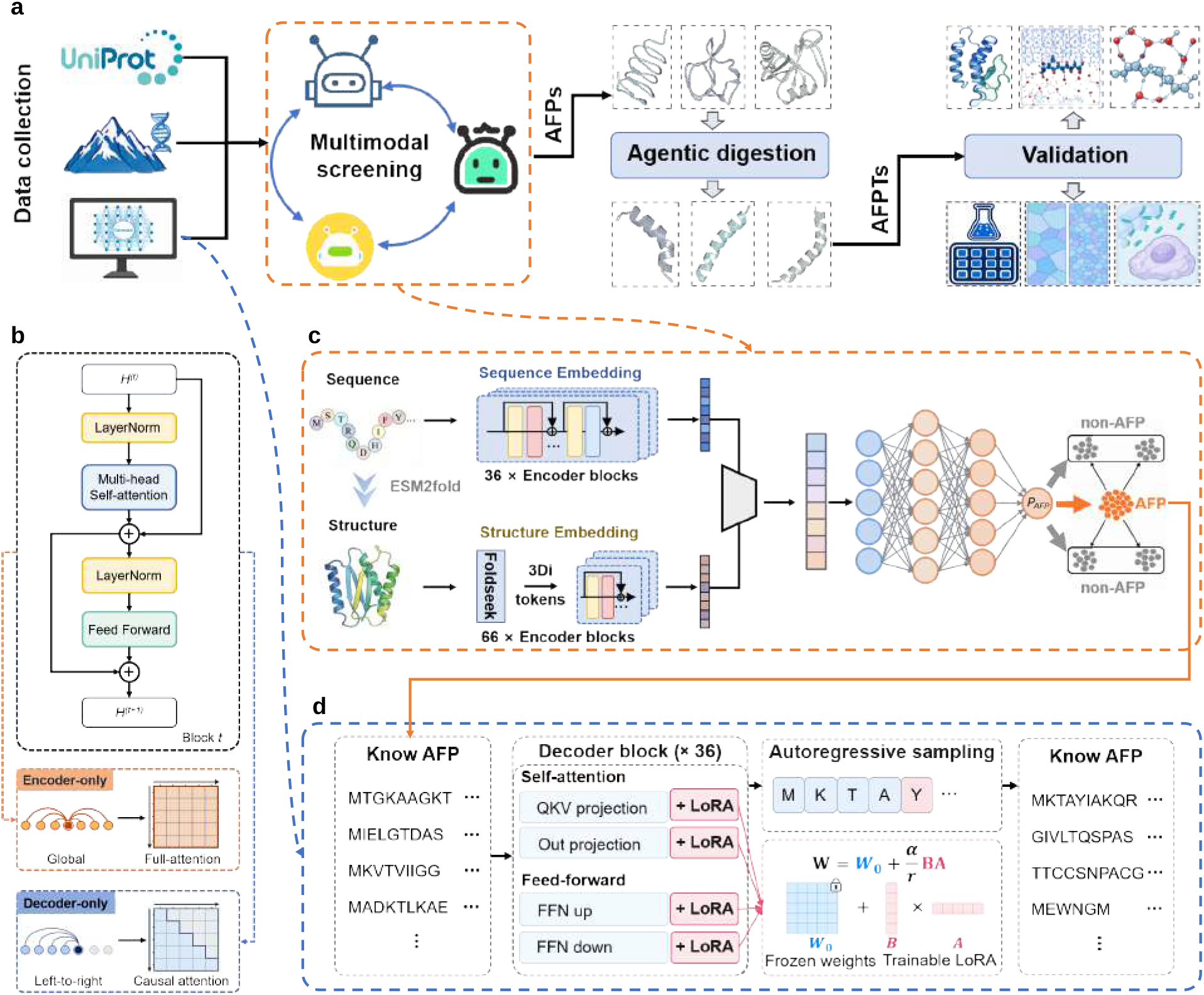
An AI-guided framework for large-scale antifreeze peptide discovery. **a**, Overview of the study workflow. Protein sequences from UniProt, glacier metagenomes and generative models are integrated and screened using multiple predictive models to identify potential antifreeze proteins. Local sequence analysis and agent-assisted candidate prioritization then identify short-peptide candidates for structural modelling, molecular simulations and experimental evaluation. **b**, Schematic of encoder-type and decoder-type Transformer architectures, illustrating bidirectional self-attention for sequence representation learning and causal self-attention for autoregressive generation, respectively. **c**, Architecture of the antifreeze protein predictor AFPFusionLM. ESM2 and SaProt extract protein sequence and structure-aware representations, respectively, which are combined through feature fusion and a multilayer perceptron for antifreeze protein classification. **d**, Domain fine-tuning of ProtGPT2 using low-rank adaptation (LoRA). Pretrained model parameters remain frozen, while trainable low-rank updates are introduced into projection layers in the attention and feed-forward networks. Candidate protein sequences are generated by autoregressive sampling.

The framework is built around complementary predictive and generative models. We developed AFPFu-sionLM, a multimodal antifreeze protein predictor that integrates ESM2 sequence representations with structure-aware representations from SaProt to recognize antifreeze-associated properties at the protein level (Fig. 1b,c). In parallel, we used low-rank adaptation (LoRA) to fine-tune the pretrained protein language model ProtGPT2 for the antifreeze domain, enabling it to produce protein candidates with AFP-associated sequence features (Fig. 1d). The predictor supports systematic screening of natural sequence resources, whereas the generator extends the search beyond existing database entries. Together, they provide the computational basis for large-scale sequence exploration.

We then connected protein-level prediction to local sequence evidence through clustering, short-fragment pattern mining and candidate prioritization, identifying peptide fragments for further investigation. Unlike candidate design based on a small number of known templates, the framework retains the relationships between local fragments, their parent proteins, related sequence groups and source databases. Candidate assessment can therefore consider both predicted function and support across sequences. This organization enables comparison of local patterns across protein backgrounds and connects large-scale computational discovery to structural simulation and experimental functional evaluation.

### Multimodal learning improves antifreeze protein prediction

To support large-scale screening across sequence sources, we systematically evaluated AFPFusionLM against six representative methods: PLM-ICE, AFP-LSE, AFProPred-AAC, CryoProtect, AFP-SRC and AFP-XGB. Evaluation covered conventional validation partitions, homology-disjoint test sets and an independent gold-standard test set, assessing classification accuracy and generalization across protein backgrounds. AFPFu-sionLM achieved leading or competitive performance in validation accuracy, recall, specificity, Matthews correlation coefficient (MCC) and area under the receiver operating characteristic curve (AUROC; Fig. 2a). Alongside the phylogenetic distribution of the evaluated sequences (Fig. 2b), homology-disjoint testing assessed recognition beyond closely related sequences. AFPFusionLM achieved AUROC and area under the precision–recall curve (AUPRC) of approximately 0.988 and MCC of approximately 0.90, exceeding the comparator methods on all three metrics (Fig. 2c). These results support the use of multimodal representations for AFP recognition under homology constraints.

**Figure 2.**
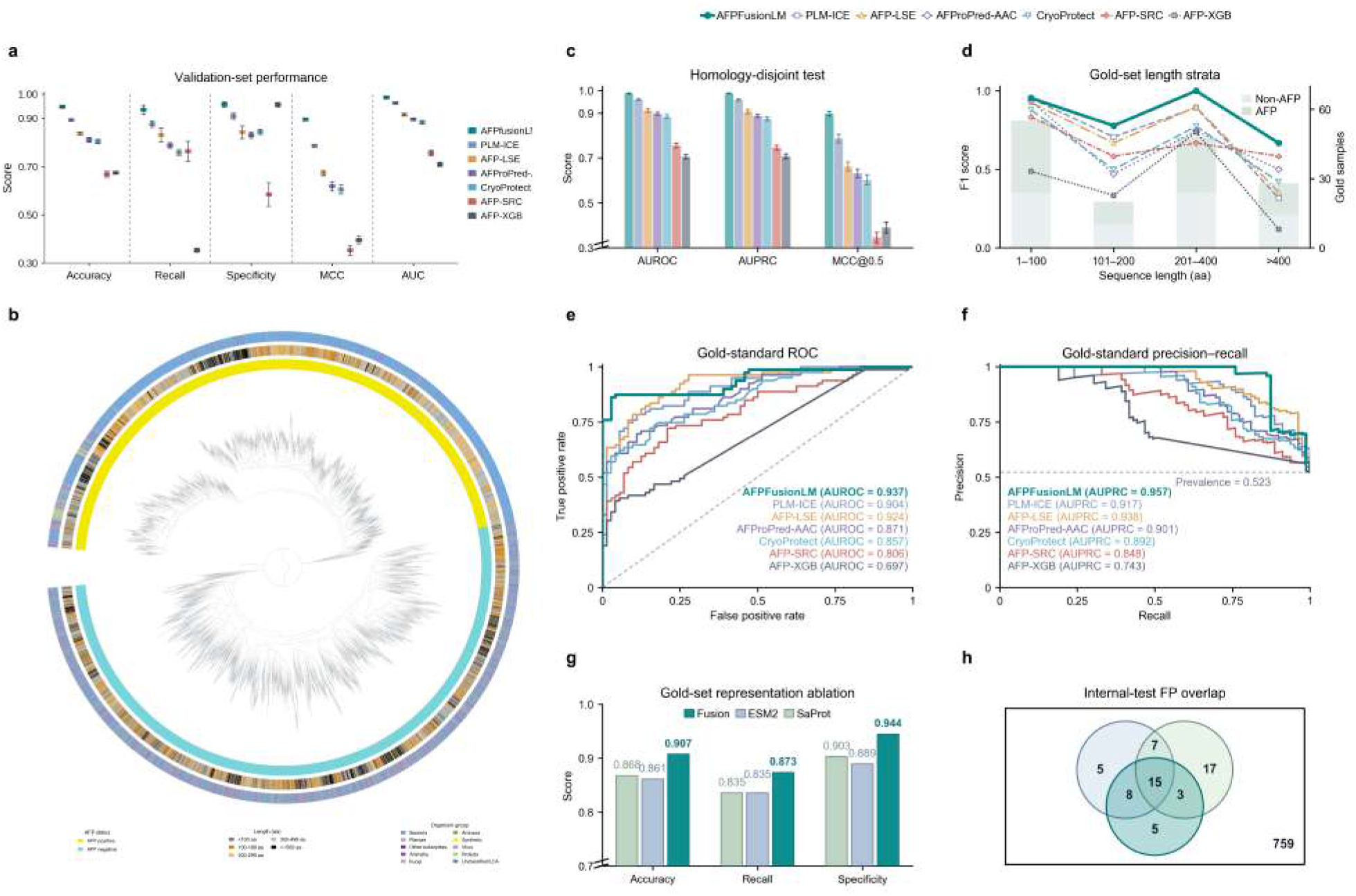
Benchmark performance evaluation and multimodal representation analysis of AFPFusionLM. **a**, Accuracy, recall, specificity, Matthews correlation coefficient (MCC) and area under the receiver operating characteristic curve (AUROC) for AFPFusionLM and six comparator methods on the randomly partitioned validation set. Points and error bars represent the mean and standard deviation across five random seeds. **b**, Circular representation of protein sequence clustering relationships in the evaluation data, with outer annotation rings indicating protein class labels and sample attributes. **c**, AUROC, area under the precision–recall curve (AUPRC) and MCC for seven methods in homology-disjoint ten-fold evaluation. Bars and error bars represent the mean and standard deviation across test folds, respectively. **d**, F1 scores for each model across four protein-length intervals in the independent gold-standard test set. Background bars indicate sample counts in the corresponding intervals. **e**,**f**, ROC and PR curves for each model on the independent gold-standard test set. AFPFusionLM achieves an AUROC of 0.937 and an AUPRC of 0.957. **g**, Ablation of sequence and structure-aware representations, comparing accuracy, recall and specificity for ESM2, SaProt and the fusion model on the gold-standard test set. **h**, Overlap of false-positive predictions from ESM2, SaProt and the fusion model on the randomly partitioned validation set. Numbers in each region indicate the corresponding numbers of protein sequences.

On the independent gold-standard test set, AFPFusionLM achieved an AUROC of 0.937, compared with 0.924 for AFP-LSE and 0.904 for PLM-ICE. Its AUPRC was 0.957, also exceeding AFP-LSE (0.938) and PLM-ICE (0.917; Fig. 2e,f). These results indicate improved overall discrimination between AFPs and non-AFPs, with high precision maintained across a range of recall levels. Length-stratified analysis further showed strong classification performance across proteins of 1–100, 101–200, 201–400 and more than 400 amino acids, with the highest F1 score in the 201–400-aa group (Fig. 2d). Although sequences longer than 400 amino acids remained more challenging, performance across length strata supports screening of protein collections with heterogeneous origins and lengths.

To examine the contribution of multimodal fusion, we compared predictions based on ESM2 sequence representations, SaProt structure-aware representations and their combination. On the gold-standard test set, accuracy increased from 0.861 for ESM2 and 0.868 for SaProt to 0.907 for AFPFusionLM. Recall increased from 0.835 for either single-modality model to 0.873, while specificity increased from 0.889 and 0.903 to 0.944 (Fig. 2g). These gains support the complementarity of sequence and structure-aware representations for identifying potential AFPs. False-positive sets on the internal held-out partition overlapped only partially (Fig. 2h), indicating differences in the errors made by the models. These observations motivate the subsequent use of multimodel consensus to increase the reliability of large-scale candidate screening.

### Generative modelling expands AFP-associated sequence space

To extend the search beyond the coverage of natural databases, we used a generative protein language model to expand the space of potential AFP candidates. Starting from pretrained ProtGPT2, we compared four adaptation strategies: a frozen backbone, soft prompting, full fine-tuning and LoRA (Fig. 3a,b). Validation and internal-test negative log-likelihood (NLL) per residue were approximately 2.84 and 2.82 for the frozen model and decreased to 2.33 and 2.29 with soft prompting. LoRA further reduced these values to 2.17 and 2.12, closely matching full fine-tuning at 2.18 and 2.12. Notably, LoRA updated only approximately 11.8 million parameters, around 1.5% of the trainable parameter count for full fine-tuning, while achieving similar sequence-modelling performance. Thus, parameter-efficient adaptation enabled the pretrained model to learn the AFP-associated sequence distribution without updating the entire backbone.

**Figure 3.**
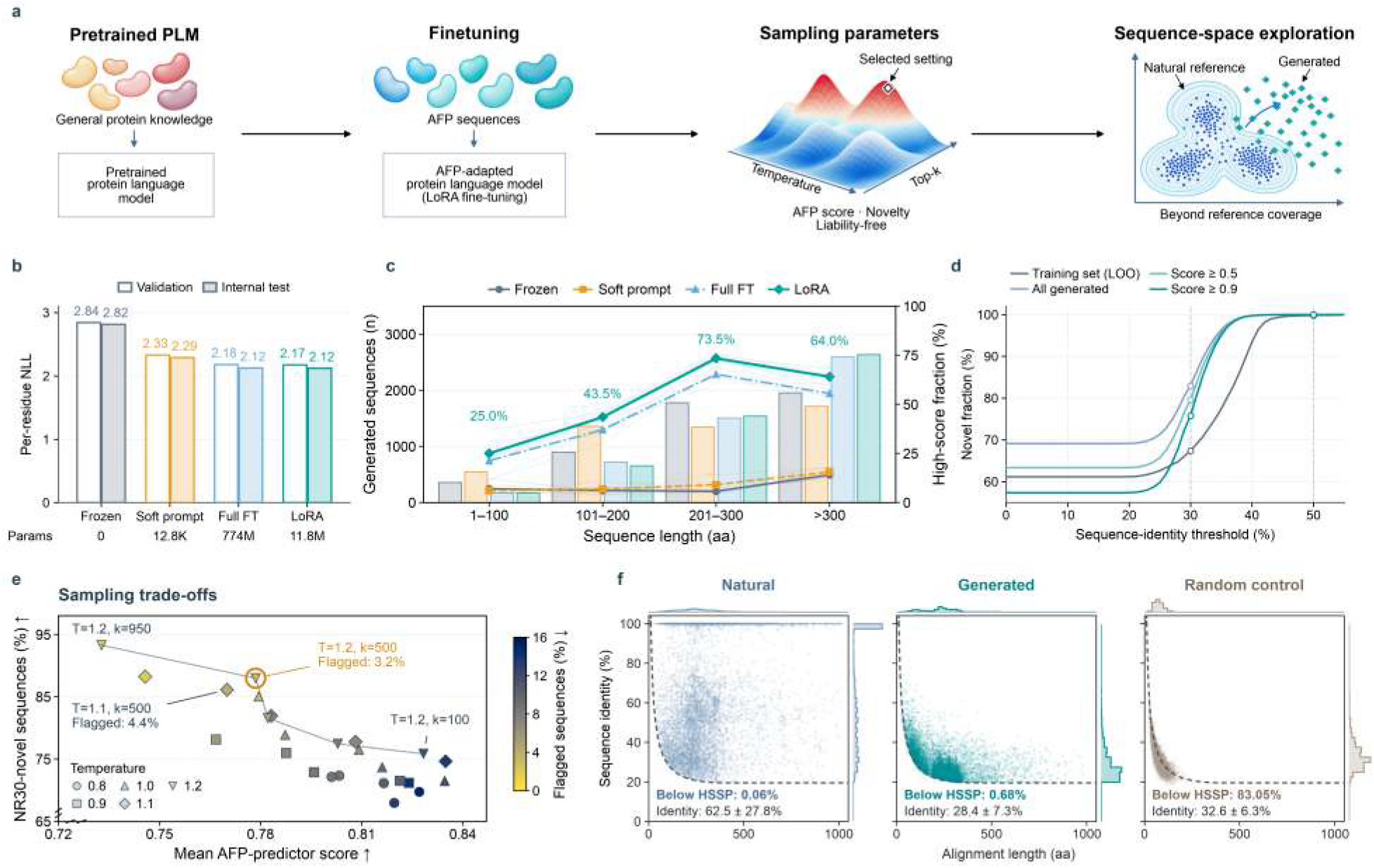
Parameter-efficient domain adaptation of ProtGPT2 and exploration of generated protein sequence space. **a**, Generative modelling workflow, comprising a pretrained protein language model, domain adaptation on antifreeze-associated sequences, sampling-parameter optimization and analysis of generated sequence space. **b**, Trainable parameter counts and negative log-likelihood (NLL) per residue on the validation and independent test sets for four adaptation strategies: a frozen backbone, soft prompting, full fine-tuning and LoRA. **c**, Numbers of sequences generated by the four adaptation strategies across protein-length intervals (bars) and fractions of sequences with an antifreeze prediction score of at least 0.9 (lines). **d**, Fractions of sequences without a qualifying training-sequence match at different sequence-identity thresholds, comparing the leave-one-out training-set reference, all generated sequences and generated subsets with prediction scores of at least 0.5 and 0.9. **e**, Relationship between mean antifreeze protein prediction score and the fraction of NR30-novel sequences across sampling-temperature and top-*k* combinations. Marker shapes distinguish sampling temperatures, and colours indicate the fraction of sequences flagged by composition- and repeat-related rules. **f**, Joint distributions of representative alignment length and sequence identity for natural proteins, generated proteins and random sequence controls aligned against a reference protein database. Marginal histograms show the corresponding variable distributions; within-panel annotations report mean sequence identity and the fraction of sequences below the HSSP threshold.

Functional prediction of generated sequences provided further support for this strategy. Under a common 120-token diagnostic sampling setting, we compared candidate yields and the proportions of high-scoring sequences across protein-length bins (score ≥ 0.9; Fig. 3c). For LoRA, the high-score fraction was approximately 25.0% for sequences of 1–100 amino acids, increased to 43.5% at 101–200 amino acids and reached 73.5% at 201–300 amino acids, remaining approximately 64.0% above 300 amino acids. This length-dependent profile shows that, within the diagnostic setting, the adapted model generated sequences with high predicted AFP potential across multiple length ranges, with the strongest enrichment among intermediate-length proteins.

The value of generative modelling also depends on its ability to explore beyond the existing sequence distribution while retaining predicted target-function characteristics. We therefore examined relationships between generated proteins and training sequences across sequence-identity thresholds, including subsets with predictor scores of at least 0.5 and 0.9 (Fig. 3d). Score-filtered sequences retained diversity and were not confined to regions of high similarity to the training data. Varying sampling temperature and top-*k* revealed a trade-off between mean AFP-predictor score and the fraction of novel sequences under the NR30 criterion (Fig. 3e). For example, temperature 1.2 and top-*k* = 500 yielded approximately 88.0% novel sequences with a mean AFP-predictor score of approximately 0.78, combining broad sequence exploration with a high predicted target-function tendency.

To further characterize generated sequences relative to natural proteins, we compared natural reference sequences, domain-adapted generated sequences and random controls using sequence alignments and the HSSP threshold (Fig. 3f). The fractions below the threshold were 0.06% for natural references, 0.68% for generated sequences and 83.05% for random controls. Under the specified representative-hit selection protocol, generated sequences thus retained stronger relationships to natural protein sequences than did random controls, while showing some distributional expansion. Subsequent functional prediction and multimodel consensus screening retained 31,948 generated protein sequences in the potential AFP collection, establishing generation as a substantial candidate source complementary to natural databases and environmental metagenomes.

### Large-scale mining reveals periodic sequence architectures

Integration of natural databases, environmental metagenomes and generated proteins produced a consensus collection of 87,305 potential AFP records: 16,746 from UniProt, 38,611 from glacier metagenomes and 31,948 from the generative model, representing 19.2%, 44.2% and 36.6% of the collection, respectively (Fig. 4a,b). The three sources showed both overlap and distinct distributions in the sequence-representation space, indicating complementary coverage of candidate sequence space. Among the UniProt candidates, 8,996 records were associated with environmental samples (53.7%), 5,215 with protists and algae (31.1%) and 2,302 with aquatic animals (13.7%). Protein-length distributions also differed: glacier-metagenome candidates were concentrated at shorter lengths, whereas generated sequences showed a high density around 200–300 amino acids (Fig. 4c). Multisource integration therefore expanded both candidate numbers and the diversity of sequence backgrounds and compositions.

**Figure 4.**
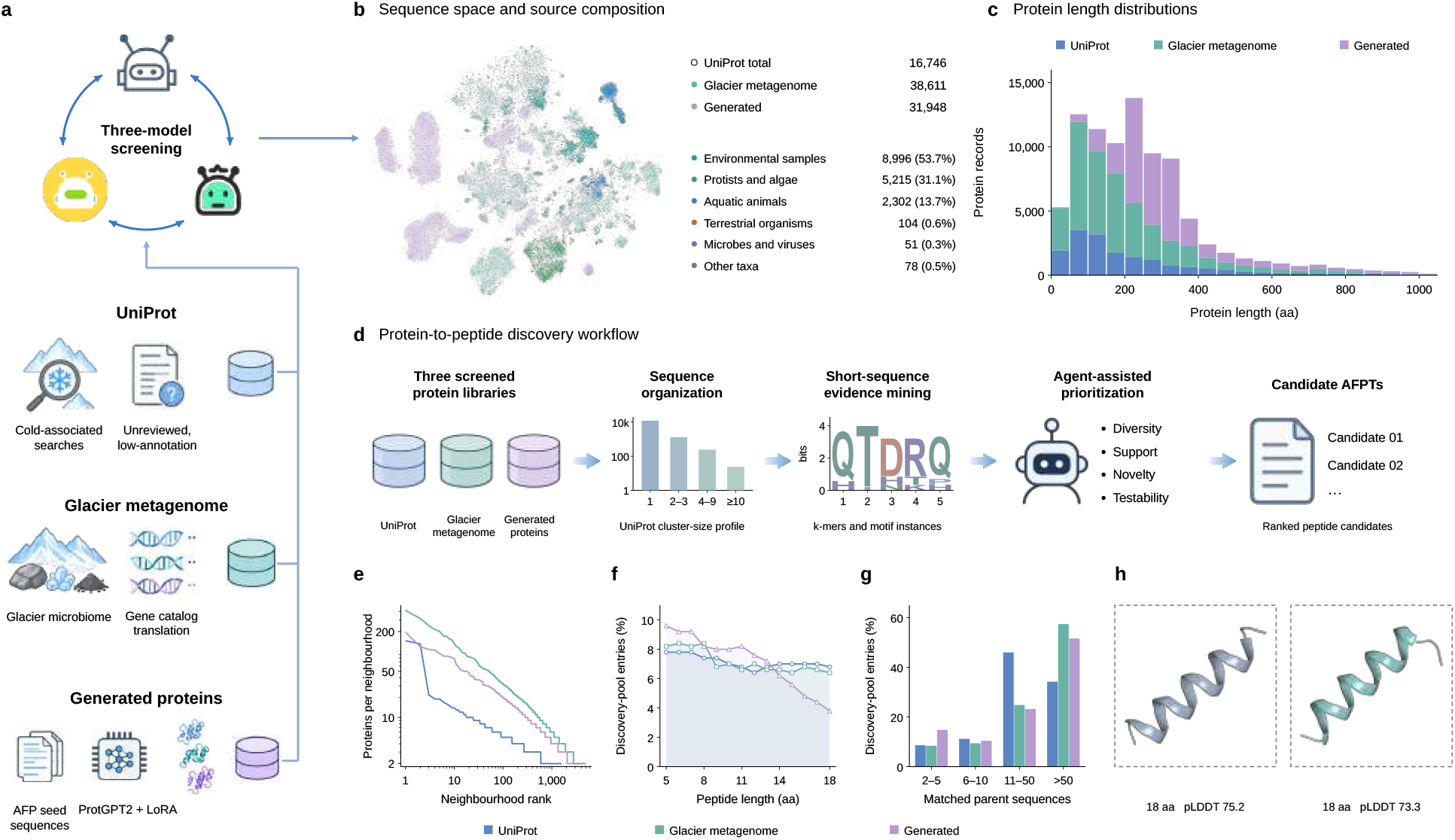
Exploration of multisource protein sequence space and prioritization of antifreeze peptide candidates. **a**, Three complementary sources of candidate proteins and a unified multimodel screening workflow: poorly annotated UniProt proteins associated with cold environments, glacier-metagenome sequences and protein sequences produced by generative models. **b**, Distribution of 87,305 candidate protein records retained by three-model consensus screening in two-dimensional sequence-representation space. The UniProt, glacier-metagenome and generated sources contain 16,746, 38,611 and 31,948 records, respectively. The source-category composition of UniProt candidate proteins is shown on the right. **c**, Sequence-length distributions of candidate proteins from the three sources. **d**, Protein-to-peptide candidate discovery workflow, comprising sequence organization, local short-sequence evidence mining and agentassisted candidate prioritization based on sequence diversity, parent-protein support, novelty and experimental feasibility. **e**, Neighbourhood-size distributions for the three protein sources, with local sequence neighbourhoods ranked by size. **f**, Fractions of entries within different peptide-length intervals in the candidate discovery pools from the three sources. **g**, Distribution of candidate-fragment support stratified by the number of matched parent proteins. **h**, Predicted conformations of two representative 18-amino-acid peptide candidates, annotated with peptide length and predicted local distance difference test (pLDDT) scores.

The large collection enabled systematic comparison of local sequence features across protein backgrounds. Under the frozen clustering parameters used for each source, neighbourhood analysis revealed different similarity profiles. Glacier-metagenome and generated sequences retained larger neighbourhoods at higher ranks, producing longer rank-size tails than UniProt (Fig. 4e). To examine potentially functional local fragments, we characterized peptide-length distributions and parent-protein support in the discovery pools (Fig. 4d,f,g). All three sources contained substantial numbers of fragments within the 5–18-aa range, although their length compositions differed. Within the archived discovery pools of 500 entries per source, entries matching more than 50 parent proteins accounted for 57.4% of the glacier-metagenome pool and 51.6% of the generated pool, compared with 34.2% of the UniProt pool (Fig. 4g). Thus, the collection contains many local features that recur across parent proteins, providing a basis for investigating sequence patterns beyond individual proteins.

More importantly, expanding the search revealed biological features beyond an increase in candidate numbers. Systematic comparison of local sequences across potential AFP clusters identified architectures with pronounced periodic organization. These architectures occurred in multiple candidate clusters, allowing local repeats previously dispersed among individual proteins to emerge as recurrent patterns across clusters. Their shared features included periodic arrangements of short sequence units and recurring natural variants, suggesting that particular residue combinations and repeat organizations may be associated with potential antifreeze function. By extending the search from a limited set of candidates to broad natural and generated sequence spaces, these features could be observed and compared across protein backgrounds, revealing potential functional sequence architectures for further investigation.

These observations guided systematic peptide discovery. We organized short-peptide candidates around periodic local fragments, considering their occurrences in parent proteins, support from related sequence groups and natural sequence variants. *k*-mer statistics and conserved-pattern analyses provided complementary evidence, while diversity, novelty and experimental tractability informed prioritization (Fig. 4d). This evidence-based organization supports comparison of repeat units, repeat numbers and local sequence changes within related candidate families. Structural prediction of two representative 18-aa candidates yielded pLDDT scores of 75.2 and 73.3 (Fig. 4h), providing reference models for subsequent structural analysis.

To further examine the effects of representative peptides on the ice–water interface, we performed 50-ns molecular dynamics simulations and analysed the development of ice-like structure using water-ordering metrics. Both peptide-containing systems showed lower ice-like water fractions than the peptide-free control. Under the *q*_6_ criterion, the time-averaged ice-like fraction decreased from 58.2% in the control to approximately 41.2–41.4% in the peptide-containing systems, with a consistent trend under the *F*_4_ criterion. Analysis of local hydration layers in the final segment of the trajectories further showed reduced orientational order and broader angular distributions near the peptides, indicating departures from a regular ice-like water network. These observations support a mechanism in which the peptides influence ice growth by perturbing the ordered arrangement of neighbouring water molecules and delaying ice-like structure formation.

In the completed ice recrystallization inhibition (IRI) screen, we evaluated 17 non-control peptide candidates using the mean area of the ten largest ice crystals normalized to the PBS control. Six candidates had relative ice-crystal areas at or below the 61.35% measured for a literature reference peptide in the same experimental batch. Notably, five of six candidates from one repeat-associated sequence family reached this reference level, with a median relative ice-crystal area of 55.88% across the family. IRI activity was therefore observed in multiple related family members rather than a single sequence. A further representative candidate produced a relative ice-crystal area of 30.26%, corresponding to a 69.74% reduction relative to PBS. These results connect sequence-level repeat patterns to measurable IRI activity, providing experimental support for discovery centred on repeat-associated families and a basis for subsequent structure–activity comparisons of repeat number, fragment boundaries and natural variants.

## Discussion

This study connects large-scale protein exploration with the discovery and functional evaluation of local peptide families. Its central contribution is a framework in which multimodal prediction and generative modelling supply candidate proteins, while local sequence analysis preserves the evidence linking short peptides to their parent sequences and related variants. Previous studies have developed antifreeze peptides by specifying ice-binding chemistry and site spacing, and have combined protein language models with molecular simulations to explore microbiome-derived candidates [8, 12]. Our study brings these lines of investigation together through multisource screening and an explicit focus on recurrent local sequence organization. This approach provides families of testable fragments through which sequence–activity relationships can be investigated.

The predictive results support the use of complementary representations for candidate screening. On the independent reference set, combining sequence and structure-aware information increased accuracy to 0.907, compared with 0.861 and 0.868 for the respective single-modality models. Performance under homology-separated evaluation further supports recognition beyond close sequence neighbours. The partial overlap among false-positive sets motivates consensus screening, which integrates outputs from three predictors that share pretrained representations. Protein-level predictions guide candidate selection, and activity is assessed through experiments on selected peptide fragments. The 87,305 retained entries form a source-labelled collection of candidate protein records for downstream sequence analysis and peptide selection.

Generative modelling provides a complementary way to explore AFP-associated sequence space. The domain-adapted generative module achieved approximately the same internal-test negative log-likelihood as full fine-tuning while updating about 1.5% as many parameters, showing that AFP-associated sequence patterns can be learned with a substantially smaller trainable parameter set. The 31,948 generated records retained after consensus screening demonstrate the contribution of this source to the candidate collection. The scoring models and generator use the same AFP-labelled sequence collection, linking generation and screening through a shared sequence basis. Within this computational workflow, prediction scores rank generated candidates for subsequent activity testing. Sequence novelty is defined relative to the specified reference collection using a sequence-identity threshold. Prospective experimental comparisons of natural and generated candidates provide a direct test of differences in functional diversity and active-candidate yield between the two sources.

Large-scale exploration revealed periodic sequence architectures recurring across multiple candidate protein clusters. This recurrence makes local sequence organization a concrete subject for functional investigation. By retaining repeat units, fragment boundaries, natural variants and parent-protein support, the workflow organizes dispersed observations into related peptide families. Experimental evaluation of peptides from this repeat-associated family confirms IRI activity and cellular cryoprotection, connecting the identified sequence patterns to measurable function. These findings support the use of repeat-associated families to guide candidate discovery and provide a basis for investigating how repeat number, local sequence variation and fragment boundaries influence activity.

Repeat-associated families also define a practical route from discovery to controlled design. Repeat number, fragment boundaries and naturally observed substitutions provide explicit variables for testing how local sequence organization affects activity. Their effects can be examined alongside composition- and lengthmatched scrambled controls, separating residue content from residue order. For subsequent family-based experiments, parent-protein relatedness, sequence composition and similarity of generated sequences to the training collection define additional comparison variables. The present evidence therefore motivates a family-based structure–activity programme in which the contribution of periodicity is tested directly. Comparisons between isolated peptides and their parent proteins could further clarify how activity depends on molecular context.

The simulations offer a molecular interpretation that can guide these experiments. Compared with the peptidefree control, peptide-containing systems showed reduced ice-like water fractions and altered local water organization. These observations are consistent with peptide-dependent changes in hydration and ice-like structure development. Earlier structural work showed that ordered surface waters can mediate binding between an AFP and ice [15]. The water-ordering metrics in the present analysis characterize collective changes in the simulated water environment. A follow-up simulation design combines independent trajectories and alternative starting configurations with spatially resolved analyses of peptide–ice contacts and hydration to examine local adsorption and broader water reorganization. These simulation results connect peptide-associated changes in water organization with ice-like structure formation, providing a molecular perspective for sequence–activity analysis.

Cell-based experiments extended the functional evidence for repeat-associated peptides to biological cryopreservation. Assessments of cytocompatibility, post-thaw survival and viability, and retained cellular function supported the cryoprotective effects of peptides from this repeat-associated family. Together with the IRI assays, these results establish functional evidence at the levels of ice recrystallization control and cellular protection during freezing and thawing. Relating the cellular outcomes to repeat units, fragment boundaries and natural variants provides an experimental basis for examining relationships between sequence architecture and biological protection. The framework thus connects multisource protein screening and local peptide discovery with molecular and cellular functional evaluation.

## Methods

### Antifreeze protein dataset construction

The supervised dataset was assembled from the AFP and non-AFP sequence collections distributed with AFProPred, retaining their source labels and supplementing the positive class with 12 traceable historical records[16]. Preprocessing comprised exact-sequence deduplication, length filtering and checks for homology to the historical reference collection. Training clusters homologous to reference sequences at 30% identity and at least 80% coverage of both query and target were excluded, and clusters containing mixed positive and negative labels were set aside. The resulting benchmark contained 15,380 distinct protein sequences: 7,193 with AFP labels and 8,187 with non-AFP labels, all shorter than 1,024 amino acids. Supervised learning data, evaluation collections and large-scale discovery libraries were maintained as separate populations with their corresponding identifiers and labels or source annotations.

Model development used label-stratified 90:10 splits, yielding 13,842 training and 1,538 validation sequences. The validation partition was used for model and checkpoint selection. For homology-separated evaluation, MMseqs2 grouped sequences into 13,518 clusters at 30% identity with at least 80% alignment coverage of both query and target[17]. Entire clusters were assigned to ten folds of 1,538 sequences without splitting clusters across folds. Rotation *k* used fold *k* for testing, the next fold for validation and the remaining eight folds for training. The historical reference collection contained 79 positive and 72 negative sequences after excluding two 2,493-residue sequences that exceeded the input range. This collection was evaluated using fixed models and was not used for parameter fitting or checkpoint selection in this study.

### AFPFusionLM architecture and training

The sequence and structure-aware branches used ESM2-3B and SaProt-1.3B-AF2 to extract 2,560- and 1,280-dimensional residue representations, respectively[18, 19]. Structure-aware inputs paired each amino acid with its Foldseek 3Di character[20]. Amino acids were retained in uppercase and 3Di characters in lowercase; the structural character was masked at positions with pLDDT below 70. Before feature extraction, amino-acid identities and lengths were checked across the input sequence, structural residues and structural alphabet, and structure-aware tokenization was required to produce no unknown tokens. Special and padding tokens were removed, and adaptive average pooling along the sequence dimension produced 16 ordered local representations.

Let *E* and *S* denote the sequence and structure-aware feature matrices, with 16 rows and 2,560 and 1,280 columns, respectively. Branch-specific projections *ϕ*_*E*_ and *ϕ*_*S*_, each comprising layer normalization, a linear transformation and GELU activation, mapped these features to 256 dimensions. Fusion was defined as

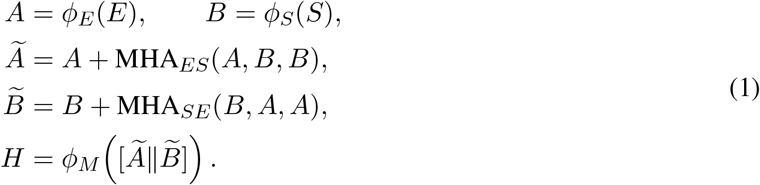

The arguments of each multihead-attention operator are the query, key and value matrices, in that order. The two directions had separate parameters, eight heads and dropout of 0.2. The symbol ∥ denotes concatenation along the feature dimension, and *ϕ*_*M*_ applies a 512-to-256-dimensional linear transformation, GELU and layer normalization, in that order. Thus, *H* contains 16 fused local representations.

For the column-vector representation *h*_*i*_ of row *i* of *H*, attention pooling and classification were

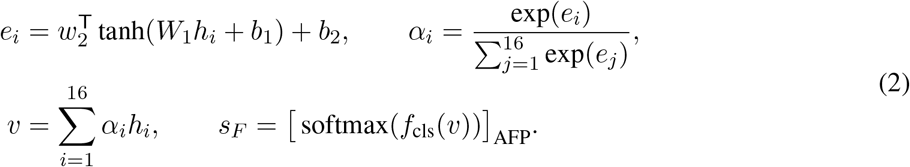

Here, *W*_1_ maps 256 features to 128 hidden units, *w*_2_ maps these units to a scalar, and *α*_*i*_ is the normalized weight of local representation *i*. The classifier *f*_cls_ comprised layer normalization, a 256-to-64-dimensional linear layer, GELU, dropout of 0.2 and a two-class output layer. The AFP score *s*_*F*_ was the positive-class softmax output. Both pretrained encoders remained frozen; the fusion and classification modules contained 1,699,523 trainable parameters.

Training used AdamW with a learning rate of 10^*−*3^, weight decay of 0.01, batch size of 128 and cross-entropy loss with label smoothing of 0.1. Runs lasted at most 60 epochs and stopped after eight epochs without validation improvement. Checkpoints were selected by validation AUROC, with AUPRC breaking ties; no learning-rate schedule was applied. Random-split experiments used seeds 17, 42, 97, 137 and 211, whereas homology-separated ten-fold evaluation used seed 42. Selected models were evaluated directly without refitting on the combined training and validation partitions.

### Model evaluation and benchmarking

Comparators were PLM-ICE, AFP-LSE, AFProPred-AAC, CryoProtect, AFP-SRC and AFP-XGB[16, 21– 25]. All methods used the same data partitions and evaluation metrics. PLM-ICE followed its public training workflow, AFP-LSE used an implementation compatible with the public architecture, and CryoProtect was retrained using its amino-acid and dipeptide composition approach. AFProPred-AAC, AFP-SRC and AFP-XGB used an amino-acid-composition adaptation, a port of the sparse-representation method and a literature-based reimplementation, respectively. Reported comparisons refer to these implementations trained under the common evaluation protocol.

AUROC and AUPRC were calculated from continuous prediction scores, with AUPRC computed as average precision. Accuracy, recall, specificity, F1 and Matthews correlation coefficient used a classification threshold of 0.5. Each protein received one out-of-fold prediction in the ten-fold analysis; the principal summaries were the mean and sample standard deviation of the test-fold metrics. ROC and precision–recall curves for the historical reference collection used the arithmetic mean of the five model scores for each sequence. Length-stratified evaluation and representation ablation used the seed-42 models. Length bins were 1–100, 101–200, 201–400 and more than 400 amino acids. Sequence-only, structure-aware and fusion models were compared on identical evaluation sequences. False-positive overlaps were calculated from negative-sequence identifiers classified as positive in the seed-42 validation partition; region counts represented intersections or differences of these error sets.

### Generative model adaptation

Generation used a pretrained autoregressive protein language model with 36 decoder layers, a hidden dimension of 1,280, 20 attention heads and approximately 774 million parameters[13]. The 7,193 AFP-labelled sequences were partitioned by NR30 homology cluster into 5,753 training, 720 validation and 720 test sequences, with no cluster shared between partitions. Sequences were tokenized by byte-pair encoding, with a training limit of 512 tokens and positions reserved for the initial and terminal tokens. Padding positions were excluded from the loss, whereas genuine terminal tokens remained supervised targets.

Four adaptation strategies were compared: a frozen backbone, soft prompting, full fine-tuning and LoRA. Soft prompting optimized ten prompt vectors, comprising 12,800 parameters. LoRA introduced low-rank updates into the joint query–key–value and attention-output projections and both feedforward projections[26]. For a projection written in output-by-input matrix convention, its effective weight at inference was

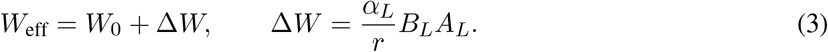

The pretrained matrix *W*_0_ has dimensions *d*_out_ ×*d*_in_, whereas *A*_*L*_ and *B*_*L*_ have dimensions *r* ×*d*_in_ and *d*_out_ ×*r*. Rank *r* was 16 and scaling coefficient *α*_*L*_ was 32. During training, dropout of 0.05 was applied to the adapter input. Backbone weights and biases were fixed, and only adapter matrices were optimized, leaving 11,796,480 trainable parameters. Learning rates selected on the validation set were 5×10^*−*5^ for full fine-tuning, 5×10^*−*4^ for LoRA and 10^*−*2^ for soft prompting.

Checkpoint selection used corpus-level negative log likelihood per amino-acid residue rather than the unweighted mean token loss:

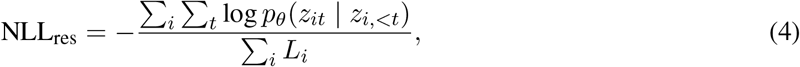

where *z*_*it*_ is target token *t* of sequence *i*, the numerator includes the log probability of the genuine terminal token, and *L*_*i*_ is the amino-acid length of the decoded protein segment. The three trainable adaptation strategies used the same five seeds used for predictor development. The held-out test partition was used to evaluate the selected checkpoints.

Before library generation, five LoRA models were refitted on all 7,193 sequences. In seed order, epoch budgets were fixed at 6, 5, 6, 9 and 5, as selected during validation. Refitting used AdamW with a learning rate of 5 ×10^*−*4^, weight decay of 0.01, a 5% warmup and linear learning-rate decay. Micro-batches contained four sequences, with gradient accumulation over eight steps, giving an effective batch size of 32. The gradient norm was clipped at 1.0, and training used BF16 precision.

### Sequence generation and novelty assessment

The four-method generation comparison used a common limit of 120 new tokens, temperature 1.0, top-*k* = 950, top-*p* = 1.0 and repetition penalty 1.2. Each method generated 1,000 outputs per seed, with paired training and sampling seeds for adapted models. Generation was initiated with one start token and stopped at the terminal token or token limit. Outputs were grouped by their decoded amino-acid lengths. High-score fractions used the scorable sequences in each length bin as the denominator and a score threshold of 0.9. Functional scores were the mean outputs of five fixed ESM2-only predictors.

Production sampling was developed using the refitted models. A seed-42 grid combined temperatures of 0.8, 0.9, 1.0, 1.1 and 1.2 with top-*k* values of 100, 200, 300, 500 and 950, generating 1,000 sequences per setting. Top-*p* was fixed at 1.0, the repetition penalty at 1.1 and the new-token limit at 512. Grid development considered five objectives: maximizing mean predictor score, NR30 novelty and NR50 self-cluster ratio, while minimizing length Wasserstein distance and amino-acid-composition Jensen–Shannon divergence. Six settings were then evaluated across five paired seeds, with minimization of sequence-liability fraction added as a sixth objective. Each confirmation run was required to achieve at least 99% terminal-token completion and 99.5% canonical amino-acid sequences, with no more than 10% exact duplicates and no more than 10% of lengths outside the training-set 0.5th–99.5th percentile interval.

A setting was Pareto-dominated if another eligible setting was no worse on every objective and strictly better on at least one. Selection within the eligible Pareto set P used the equally weighted geometric mean of directionally aligned percentile ranks:

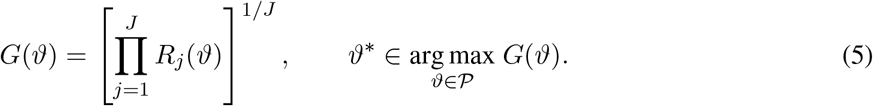

Here, *ϑ* denotes a temperature–top-*k* setting and *R*_*j*_ its normalized percentile rank for objective *j*, oriented so that larger values are preferable. The number of objectives *J* was five during grid development and six during confirmation. Ties in the final composite score were resolved by lexicographic order of the setting identifier. This score selected sampling settings, not individual proteins or peptides. The 50,000-sequence production library used temperature 1.1 and top-*k* = 500, with 10,000 outputs from each model. The temperature-1.2, top-*k* = 500 setting shown in the figure belongs to the parameter-grid comparison.

Sequence-liability flags were defined relative to the training distribution. Features were the longest homopolymer run, longest hydrophobic run, amino-acid-composition entropy, maximum single-residue fraction, |*n*_K_ + *n*_R_ − *n*_D_ − *n*_E_|/*L* and hydrophobic fraction, with A, V, I, L, M, F, W and Y treated as hydrophobic. A sequence was flagged when either run length, the maximum single-residue fraction or the absolute charge-composition measure exceeded the training-set 99.5th percentile, entropy fell below the 0.5th percentile, or hydrophobic fraction lay outside the 0.5th–99.5th percentile interval. The flagged fraction summarized composition and contiguous-repeat characteristics of the generated sequences.

Novelty relative to the training collection was assessed by MMseqs2, requiring at least 80% coverage of both query and target. Let *Q* be the eligible query collection for an analysis, *Q*_uniq_ its exact-sequence-deduplicated set and M_*τ*_ (*q*) the reference hits for query *q* meeting identity threshold *τ* and both coverage requirements. Reference novelty and internal diversity were defined separately as

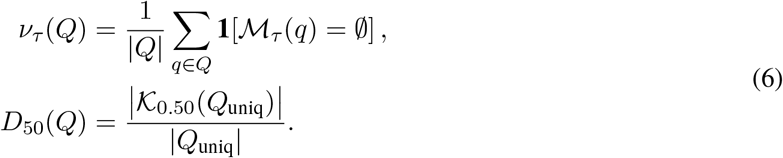

The indicator **1**[·] equals one when its condition holds and zero otherwise. NR30 corresponds to *v*_0.30_; *K*_0.50_ is the set of self-clusters at 50% identity and at least 80% bidirectional coverage. Fractions were multiplied by 100 when reported as percentages. Queries without qualifying hits remained in the novelty denominator. Sampling-grid novelty and self-clustering used distinct canonical sequences. Novelty curves for the complete generated collection and subsets scoring at least 0.5 or 0.9 were summarized separately from their respective query lists. Searches used sensitivity 5.7, an *E*-value threshold of 10^*−*3^ and at most 50 targets, followed by verification of bidirectional coverage. Leave-one-out comparisons excluded the training query itself. Reference novelty measures absence of qualifying reference matches, whereas *D*_50_ measures diversity within the generated collection.

Comparison with public protein space used 10,000 natural-source candidates, 10,000 generated candidates and 10,000 random controls. Natural and generated sequences were matched without replacement in 10-residue length bins using seed 42. Each random control retained the length of its corresponding generated sequence and was sampled independently and uniformly from the 20 standard amino acids plus X, B, U, O and Z. HHblits 3.3.0 searched UniRef30 2023_02 for two iterations, using an inclusion *E*-value of 10^*−*3^, a reporting probability threshold of 20%, a reporting *E*-value limit of 10^6^ and at most 500 reported hits[27]. For alignment length *L*, the HSSP identity threshold was implemented as

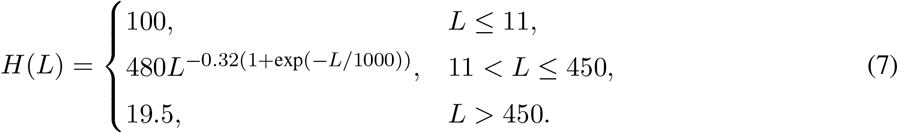

One representative hit was selected per query. Hits at or above *H*(*L*) were prioritized by alignment length, then identity, hit probability and lower *E*-value. If none reached the threshold, selection maximized identity minus *H*(*L*) before applying the same tie-breakers. All queries contributed to the below-threshold denominator; queries without hits were included in that fraction but were not assigned scatter coordinates. The horizontal axis therefore represents representative alignment length, not full query length.

### Construction of multi-source protein libraries

The public natural-protein library comprised unreviewed, sparsely annotated UniProtKB/TrEMBL entries. The query combined cold-environment, polar or marine terms in organism names with low-annotation terms in protein names. Source terms included antarctic, arctic, polar, psychrophile, psychrotolerant, cold-adapted, alpine, glacier, sea ice, permafrost, marine, subzero, freeze-tolerant, overwintering and ice-associated. Annotation terms included uncharacterized, hypothetical, predicted, putative, unnamed protein product, unknown, function unknown, conserved hypothetical and DUF. Terms were joined by OR within each set and by AND between sets. The April 2026 export contained 623,468 records. Excluding 5,194 sequences longer than 1,022 residues left 618,274 records for sequence scoring. Source mappings were retained without similarity-based dereplication, and unknown residues retained their original symbols.

The glacier library used the published Tibetan glacier microbiome gene catalogue[28], which covers 85 metagenomes and 883 cultured isolates from 21 glaciers, including ice, snow and cryoconite environments. Published translations of predicted open reading frames were used directly, without reassembling metagenomes. The downloaded protein file contained 25,320,680 records. Records containing X or having raw sequence strings longer than 1,022 characters were removed, after which terminal stop characters were stripped, yielding 25,162,254 sequences for scoring. The source project identifiers are OEP003083 and PRJNA813429.

The generated library was the five-model production collection described above. Of 50,000 outputs, 49,996 had non-empty decoded sequences, 49,905 contained canonical amino acids and 49,231 received sequence-predictor scores. Output, canonical-sequence and scorable-sequence counts were tracked separately. The 32,338 generated records with fixed five-model ESM2 mean scores greater than 0.9 entered structural evaluation. For natural sources, structural evaluation used the previously established sequence-score prescreening collections of 23,658 UniProt and 39,127 glacier candidates. These fixed collections underwent common structural processing and model reassessment.

### Consensus-based candidate screening

Candidate protein structures were predicted with ESMFold2-Fast and its ESMC-6B backbone using BF16 precision, three loops, 50 sampling steps, one diffusion sample and seed 0, without multiple-sequence alignments or structural templates. Foldseek converted the predicted structures to 3Di representations, which were processed using the pLDDT masking and case conventions described above. Amino-acid identities and lengths were checked across sequences, structural residues and structural characters. Consensus decisions were restricted to records passing these checks and receiving valid scores from all three models.

Let *s*_*E*_, *s*_*S*_ and *s*_*F*_ denote the ESM2, SaProt and fusion scores, respectively. Records were retained according to

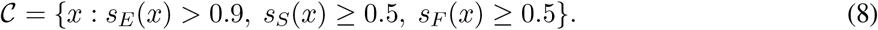

For UniProt and glacier sources, *s*_*E*_ was the seed-42 ESM2-only score; for generated sequences, it was the mean of the five ESM2 models. SaProt and fusion scoring both used fixed seed-42 models. The branches used separately fitted classifiers while retaining the pretrained representations shared by their respective architectures. The retained collection contained 16,746 UniProt, 38,611 glacier and 31,948 generated records. The total of 87,305 denotes source-labelled protein records; records with identical sequences retained their individual source identities.

Sequence-space visualization used mean-residue ESM2 representations. Embeddings for the 95,123 prescreened records were jointly standardized and reduced to 50 principal components before two-dimensional multiscale embedding with openTSNE 1.0.4. Settings were perplexities of 50 and 500, Euclidean distance, principal-component initialization, early exaggeration of 12 for 250 iterations, 500 subsequent iterations and seed 42. Source labels were not used in fitting. The 87,305 final candidates were matched to the fixed coordinates by identifier and sequence identity. UniProt source categories were assigned from the original organism or environmental-sample annotations; length distributions and source proportions used the corresponding candidate-record totals as denominators.

### Protein-to-peptide discovery and prioritization

Peptide discovery targets observed local protein segments, prioritizing tandem repeats and their natural variants while using shared *k*-mers and statistical motifs as complementary evidence. Identical full-length sequences form one sequence-support unit, with all original records and source mappings retained. Unknown or non-standard residues delimit analyzable stretches of canonical amino acids, and searches do not span these boundaries. Proteins are clustered separately by source, while small and singleton clusters remain within the scope of repeat scanning.

Local sequence groups are organized by MMseqs2 greedy length-based clustering. Identity thresholds of 30%, 40% and 50% are compared at 60% target-sequence coverage, using coverage mode 1 and cluster mode 2. Threshold selection considers cluster-size distributions, singleton fractions, protein coverage by clusters of at least five and at least ten distinct sequences, within-cluster length variation and representative-to-member alignments, without tuning for the recovery of a specified peptide. The three-source neighbourhood analysis used identity thresholds of 40% for UniProt, 40% for glacier sequences and 30% for generated sequences. Neighbourhood sizes are ranked by member-record count; motif support is counted using distinct full-length sequences.

Tandem-repeat detection searches unit lengths of 3–60 amino acids and requires at least two complete, directly adjacent units. A candidate tract *T* is partitioned without gaps into *m* units of equal length *l*. Exact and approximate repeats satisfy

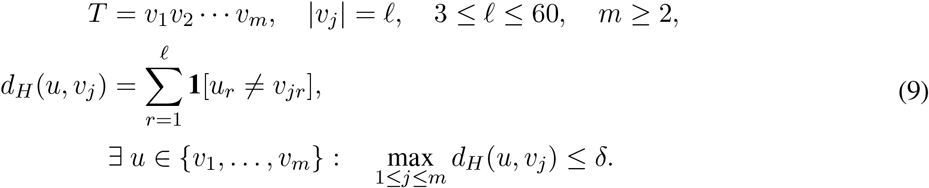

Here, *d*_*H*_ is Hamming distance, *δ* = 0 defines an exact repeat and *δ* = 1 permits at most one substitution per unit relative to a common observed anchor *u*. This criterion applies uniformly across the searched periods, without insertions, deletions or transitive relaxation through successive variants. The anchor may occur anywhere within the tract, and an exact-repeat seed is not required. The shortest fundamental period explaining the entire tract is preferred; longer harmonic periods are retained as redundant associations. Maximum single-residue fraction, *f*_max_ = max_*a*_ *n*_*a*_/*L*, is assessed separately for anchors, repeat tracts and extracted peptides, where *n*_*a*_ is the count of amino acid *a* and *L* the length of the object being assessed. Objects are excluded only when *f*_max_ *>* 0.80; equality is retained.

Within each protein cluster, cyclically equivalent units receive a common pattern identifier while retaining their observed starting phases. Reversed sequences and composition-matched permutations are treated separately. Anchors must be observed sequences; competing assignments are resolved by exact-repeat support, total support and then lexicographic order of the canonical identifier. After pattern assignment and composition filtering, let *P*_*c*_ be the set of distinct full-length sequences in cluster *c*, and *E*_*gc*_ and *V*_*gc*_ the subsets supporting pattern *g* under the exact and approximate rules, respectively. Support is calculated as

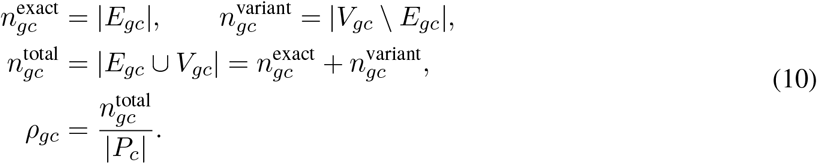

Exact support requires at least two contiguous identical units. The variant count denotes additional proteins only, preventing a protein with both exact and approximate sites from being counted twice. The fraction *ρ*_*gc*_ uses the entire distinct-sequence cluster as its denominator, not only proteins containing a detected repeat. Multiple sites within one protein increase site counts without increasing sequence support. Distinct-sequence support measures recurrence across sequences rather than independent evolutionary origins. Identical patterns across clusters are linked through an index while retaining cluster identities.

Shared-fragment discovery enumerates all contiguous 5–18-residue *k*-mers in clusters containing at least two distinct full-length sequences. Fragments are retained when supported by at least two distinct sequences and passing the composition filter. Nested fragments are assigned to the same direct-extension group only if their supporting protein sets are identical and all occurrences share a fixed positional offset. Conserved motifs are identified with MEME Suite 5.5.9 in clusters containing at least four distinct analyzable sequences[29]. ZOOPS and ANR models are run separately, with widths of 5–18 residues, up to ten motifs per run, seed 42 and a zeroth-order amino-acid background estimated from the entire distinct-sequence pool. FIMO maps motifs back to the same cluster, retaining matches with *p <* 10^*−*4^ and recording coordinates, matched segments and corresponding *q* values[30]. Shared-fragment and MEME searches operate independently of repeat detection. Statistical motif consensuses are stored separately from observed peptide sequences.

All contiguous 5–18-residue windows are extracted from retained repeat tracts and linked to shared-fragment and observed FIMO matches. Each candidate retains its parent protein, source, protein cluster, pattern group, one-based inclusive coordinates and up to ten flanking residues on either side. Identical peptide sequences share one candidate identifier while retaining all source and site mappings. A peptide is labelled tandem-repeat-containing only if it itself contains at least two complete adjacent units. Review prioritizes patterns with exact-repeat support from at least two distinct proteins, followed by multisupported variant patterns; single-protein observations and complementary discoveries are organized separately. Within pattern groups, comparisons address short cores, natural extensions, repeat numbers, extraction phases and observed single-residue changes, avoiding repeated nomination of adjacent windows as separate discoveries. Cluster-level workbooks and complete site tables preserve the evidence linking prioritization to the original sequences.

### Structural modelling and ice–water molecular dynamics

Predicted conformations of representative peptides were used to display local structure, with prediction confidence reported as pLDDT. Ice–water simulations used GROMACS 2024.6, the AMBER99SB-ILDN force field and TIP4P/Ice water[31, 32]. A single peptide was placed in a preconstructed 3.60 × 4.68 × 7.00 nm^3^ ice–water system. Overlapping waters were removed and counterions were added to ensure electroneutrality; a peptide-free system served as the reference. Conjugate-gradient minimization used at most 10,000 steps and a maximum-force convergence criterion of 100 kJ mol^*−*1^ nm^*−*1^. The integration step was 2 fs, with V-rescale temperature coupling and semi-isotropic Berendsen pressure coupling. Periodic boundaries were applied in all dimensions. Short-range electrostatic and van der Waals cutoffs were 1.2 nm, long-range electrostatics used particle-mesh Ewald, bonds involving hydrogen were constrained and coordinates were saved every 20 ps.

Water ordering was characterized using local *q*_6_ and *F*_4_. The sixth-order spherical-harmonic parameter *q*_6_ used the four nearest oxygen neighbours of each water molecule, with *q*_6_ *>* 0.55 classified as ice-like. For *F*_4_, neighbouring waters within 0.35 nm were used to construct H–O–O–H dihedrals from the most widely separated hydrogen pair. The neighbourhood mean of cos(3*ϕ*) was calculated, with *F*_4_ *<* −0.3 classified as ice-like. The fraction satisfying each criterion was calculated per frame and summarized over 0–50 ns. Local hydration analysis used waters within 0.35 nm of the peptide during the final 5 ns.

### Experimental evaluation and statistical analysis

IRI evaluation used representative peptides synthesized during the earlier candidate-discovery stage, with peptide-free PBS and literature reference peptides as comparators. Sample droplets were released from a height of 1.4 m onto a precooled substrate to form thin polycrystalline ice, annealed at −6 ^*°*^C and imaged by optical microscopy. ImageJ measured the ten largest crystals in each analyzed field[33]. Their mean area was normalized to the corresponding PBS reference:

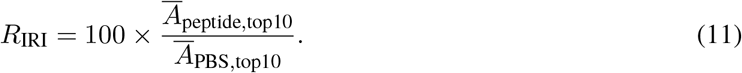

*R*_IRI_ is the percentage of crystal area remaining relative to the control, with lower values indicating stronger recrystallization inhibition under the measurement conditions; it is not the percentage inhibition itself. Candidates, concentrations, measurement groups and controls were linked to their experimental records, and missing measurements were not replaced with zero. The ten crystals contributed to one field-level mean; candidate comparisons used descriptive summaries of the available measurements.

Random-split predictor experiments were summarized by means and sample standard deviations across five seed-specific runs, and homology-separated evaluation across ten test folds. For generation, the repetition unit was the paired training and sampling seed; deterministic frozen-backbone NLL was calculated separately. High-score fractions and generation-completion rates used scorable sequences and generation attempts as their respective denominators. Novelty fractions used all eligible query sequences, including queries with no qualifying match. HSSP distributions contained 10,000 queries per group, with alignment length and identity reported as mean and sample standard deviation. Repeat-pattern support was counted by distinct full-length protein sequence. MEME/FIMO statistics were interpreted within their respective motif and search scopes; *q* values were not pooled as a global false-discovery rate across protein clusters. Crystals within an IRI field, water molecules and frames from the same trajectory were treated as subsamples of their corresponding measurements. IRI and interfacial simulations were compared descriptively without additional between-group significance tests.

## Data availability

Project GitHub repository: https://github.com/Knitua/AFPfusionLM.

## Author contributions

Yijun Li developed the models, performed the experiments, wrote the manuscript and prepared the figures. Chi Wang developed the models, wrote the manuscript and prepared the figures. Jialin Liu performed the experiments and wrote the manuscript. Mingye Song, Letian Zhang and Shuzhe Sun provided experimental guidance. Wentao Li, Jiangjie Qiu and Yihua Qin provided guidance on model development. Jinhao Yao provided experimental guidance and performed validation. Leyi Zhao performed validation. Huimin Yu and Xiaonan Wang supervised and reviewed the overall study. Yijun Li, Chi Wang and Jialin Liu contributed equally to this work.

## Additional information

